# PEELing: an integrated and user-centric platform for spatially-resolved proteomics data analysis

**DOI:** 10.1101/2023.04.21.537871

**Authors:** Xi Peng, Jody Clements, Zuzhi Jiang, Shuo Han, Stephan Preibisch, Jiefu Li

## Abstract

**Summary:** Molecular compartmentalization is vital for cellular physiology. Spatially-resolved proteomics allows biologists to survey protein composition and dynamics with subcellular resolution. Here we present PEELing, an integrated package and user-friendly web service for analyzing spatially-resolved proteomics data. PEELing assesses data quality using curated or user-defined references, performs cutoff analysis to remove contaminants, connects to databases for functional annotation, and generates data visualizations—providing a streamlined and reproducible workflow to explore spatially-resolved proteomics data.

**Availability and Implementation:** PEELing and its tutorial are publicly available at https://peeling.janelia.org/. A Python package of PEELing is available at https://github.com/JaneliaSciComp/peeling/.

**Contact:** Technical support for PEELing.

## Introduction

Localization and function of proteins are always coupled. For instance, proteins for intercellular adhesion and communication are localized to the cell surface while many energy-producing enzymes stay in the mitochondrion. High-resolution, proteome-wide mapping of protein localization is of core importance for understanding cellular organization and processes. Emerging technologies for spatially-resolved proteomics, particularly by proximity labeling, make this possible and broadly applicable in cell biology (Roux *et al.*, 2012; Rhee *et al.*, 2013; Branon *et al.*, 2018; Loh *et al.*, 2016; Geri *et al.*, 2020; Han *et al.*, 2018; Qin *et al.*, 2021). Like other enrichment-based profiling methods, spatially-resolved proteomics can be interfered by ineffective enrichment, non-specific contamination, and other factors. Rigorous assessment of data quality and proper data processing is crucial for interpreting the results and for designing subsequent studies. However, this can be complex and overwhelming, particularly for biologists with limited proteomics experience or systems biology background. To address this, we built PEELing (*p*roteome *e*xtraction from *e*nzymatic *l*abel*ing* data), a platform integrating data quality checks, contaminant removal, functional annotation, and visualization into an automated workflow (**Figure 1a**).

**Figure 1.**
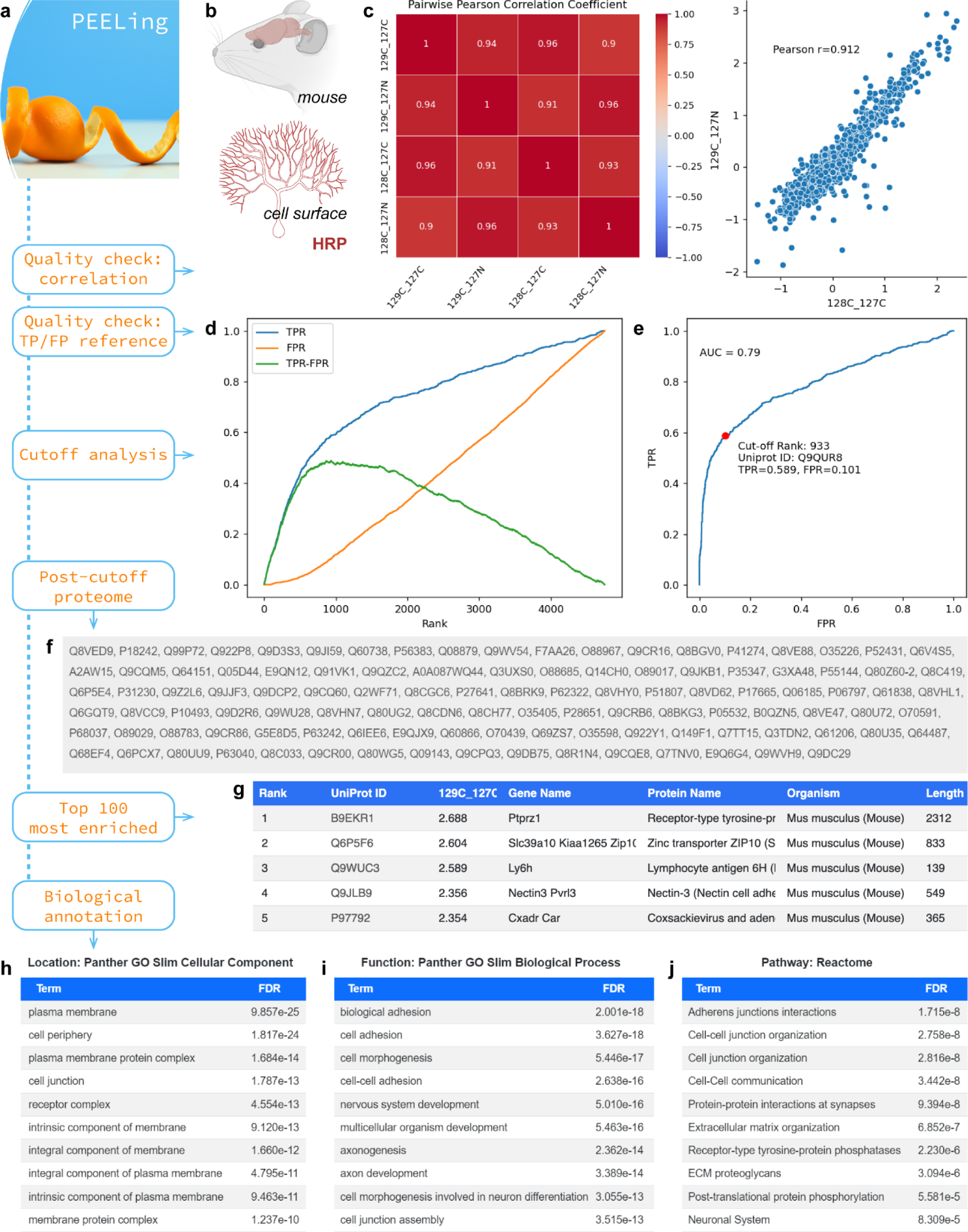
PEELing workflow and the analysis of a mouse cell-surface proteome. (**a**) PEELing workflow. (**b**) Shuster *et al.*(Shuster *et al.*, 2022) used horseradish peroxidase (HRP) mediated cell-surface biotinylation to capture the cell-surface proteome of mouse Purkinje cells at postnatal day 15. The authors used tandem mass tag (TMT) (Thompson *et al.*, 2003) based quantitative mass spectrometry. 129C and 128C TMT tags were used for cell-surface-labelled samples while 127C and 127N tags were used for non-labelled controls, yielding four labelled-to- control ratios (**Supplementary File 1**). (**c**) Correlation plots and coefficients. (**d**) True positive rate (TPR, blue), false positive rate (FPR, orange), and their difference (TPR–FPR, green) plotted against 129C:127N ratio-based ranking (x-axis). (**e**) Receiver operating characteristic (ROC) curve, based on ranking by 129C:127N. Red dot, cutoff position. AUC, area under the curve. (**f**) Post- cutoff proteome. (**g**) Top 5 most enriched proteins based on 129C:127C. On the website, this list extends to the top 100. Length unit: amino acids. (**h–j**) Protein ontology analyses for localization (**h**), function (**i**), and pathway (**j**). FDR, false discovery rate. The PEELing icon was produced by DALL-E of OpenAI.

## Implementation, Benchmarks, and Results

In **Figure 1**, we used a published cell-surface proteome of mouse developing Purkinje cells (Shuster *et al.*, 2022) (**Figure 1b** and **Supplementary File 1**) to demonstrate the functionalities of PEELing. In this study (Shuster *et al.*, 2022), cell-surface proteins were chemically labelled and isolated (“labelled” groups, hereafter) using horseradish peroxidase (HRP) mediated cell-surface biotinylation (Loh *et al.*, 2016; Li *et al.*, 2020; Shuster *et al.*, 2022). Non-labelled control groups (“control”) were included to capture contaminants such as endogenously biotinylated proteins and non-specific binders to isolation reagents. PEELing uses a ratiometric strategy as previously described (Hung *et al.*, 2014) in which the labelled-to-control ratio of each protein reflects whether this protein is cell-surface enriched or not. A *bona fide* cell-surface protein should exhibit a high ratio because it should be enriched in the labelled group relative to the control group. A contaminant should have a low ratio since it should be captured similarly in both labelled and control groups.

To assess data quality, PEELing started with pairwise correlation analysis to check whether biological replicates exhibited consistency (**Figure 1c**). To determine whether cell-surface proteins were enriched, PEELing ranked all detected proteins in descending order based on the labelled-to-control ratio and scanned through them to mark curated cell-surface proteins (true positives, TPs; **Supplementary File 10**) and intracellular contaminants (false positives, FPs; **Supplementary File 12**). As shown in **Figure 1d**, true positive rate (TPR, blue), false positive rate (FPR, orange), and their difference (TPR–FPR, green) were calculated and plotted against ratio-based ranking. TPR increased quickly while FPR rose slowly, which led to a single peak of TPR–FPR, revealing a high ranking and an effective enrichment of cell-surface proteins. This is further illustrated by a receiver operating characteristic (ROC) curve bending towards the left-upper corner and exhibiting an area under the curve (AUC) of 0.79 (**Figure 1e**).

To test whether PEELing detects data of poor quality, we generated a pseudo dataset with random values mimicking a failed enrichment (**Supplementary File 2** and **Supplementary Figure 1**). TPR, FPR, and ROC curves all followed the diagonal line (**Supplementary Figure 1b,c,e,f**), showing a complete mix of cell-surface and intracellular proteins without any enrichment (**Supplementary Figure 1d,g**). We then tested reference specificity by analyzing the cell-surface proteome (Shuster *et al.*, 2022) (**Supplementary File 1**) with nuclear and mitochondrial references (**Supplementary File 14–17**). No enrichment of proteins of unintended cellular compartments (**Supplementary Figure 2**) showed the quality of the example data as well as the necessity of data-reference matching in PEELing analysis.

All analyses above used the labelled-to-control ratio as the enrichment indicator. Protein abundance is widely used in proteomic data quantification; however, it is not a reliable enrichment indicator in spatially-resolved profiling compared with the labelled-to-control ratio (**Supplementary Figure 3**, compared with **Figure 1**). From the same cell-surface proteome dataset (Shuster *et al.*, 2022), we obtained protein abundance values of the labelled groups (**Supplementary File 3**). Despite comparable correlations based on abundance (**Supplementary Figure 3a**) and labelled-to-control ratio (**Figure 1c**), cell-surface protein enrichment was not observed when ranking the proteome by abundance (**Supplementary Figure 3b,c,e,f**). Intracellular proteins were highly enriched instead (**Supplementary Figure 3d,g**). Notably, a contaminant (e.g., a non-specific binder to the isolation reagent) can be abundant while a cell-surface protein may have a low expression level. As long as the control group captures the contaminant but not the cell-surface protein, these proteins will be ranked correctly by the labelled-to-control ratio instead of being ranked reversely by abundance. Therefore, labelled-to-control ratio is the preferred input data for PEELing.

Following data quality checks, PEELing performed cutoff analysis to remove contaminants. For each labelled-to-control ratio, PEELing found the ranking position where TPR–FPR was maximal, as indicated by the peak of the green line in **Figure 1d** and the red dot in **Figure 1e**, and retained proteins ranked above this position. The “TPR–FPR maximum” cutoff provides two key benefits: 1) The cutoff position is determined by data quality rather than an arbitrary value, allowing for unbiased assessment of the proteomic results. If the data is of high quality with sparse false positives, TPR–FPR will peak later in the ranking, resulting in the retention of more proteins.

If the data is heavily contaminated with false positives, TPR–FPR will peak earlier, leading to the retention of fewer proteins. 2) Any protein ranked above the cutoff position is retained, regardless of how it is annotated by a database. Therefore, missing annotations or occasional incorrect annotations in databases will not impact the analysis, as long as they are largely accurate and comprehensive. As illustrated in **Supplementary Figure 4**, 10-fold reduction of reference coverage, in either TP/FP or both, did not impair the analysis of the cell-surface proteome of Purkinje cells (compared with **Figure 1**). Additionally, the *Drosophila* proteome has less complete annotations than those of mouse and human, leading to jagged TPR, FPR, and ROC curves (**Supplementary Figure 5c,d**). Nevertheless, it did not interfere with the analysis of a *Drosophila* neuronal surface proteome (Li *et al.*, 2020) (**Supplementary Figure 5** and **Supplementary File 4**). Importantly, in certain physiological or pathological contexts, intracellular proteins can be transported to the cell surface—for instance, RNA helicase U5 snRNP200 in acute myeloid leukemia (Knorr *et al.*, 2023). PEELing retains these proteins if they are ranked highly, potentially enabling researchers to discover novel biomarkers and cellular processes. Despite its robustness to varied reference coverages, we note that PEELing relies on high-quality references (see Supplementary Materials – Method Details on how to create references). When references are not possible to obtain, commonly used statistical analysis is an alternative approach.

PEELing conducted cutoff analysis on all submitted labelled-to-control ratios individually and, for the final proteome, retained only those proteins that passed the cutoff of all ratios, which further removed contaminants. As shown in **Figure 1f**, PEELing displayed the post-cutoff proteome and provided information of the top 100 most enriched proteins for each labelled-to-control ratio (**Figure 1g**). On the website, each UniProt accession number is a clickable link to the corresponding UniProt protein page (Consortium *et al.*, 2023). PEELing then transmitted the post-cutoff proteome to the Panther server (Mi *et al.*, 2019; Thomas *et al.*, 2022) for over-representation analyses on protein localization (**Figure 1h**), function (**Figure 1i**), and pathway (**Figure 1j**) and revealed an enrichment of cell-surface proteins related to cell adhesion and neuronal development, perfectly matching the dataset—a cell-surface proteome of developing Purkinje cells.

Simply plugging in corresponding references, PEELing is ready for analyzing spatially-resolved proteomics data of any cellular compartments, such as the nucleus (**Supplementary Figure 6**) by TurboID (Branon *et al.*, 2018) (**Supplementary File 5**) and miniTurbo (Branon *et al.*, 2018) (**Supplementary File 6**), as well as the mitochondrion (**Supplementary** Figure 7 and 8) by APEX2 (Lam *et al.*, 2014; Han *et al.*, 2017) (**Supplementary File 7**) and BioID (Roux *et al.*, 2012; Branon *et al.*, 2018; Antonicka *et al.*, 2020) (**Supplementary File 8** and **9**). Together, we demonstrated that PEELing is widely applicable to: 1) various organisms, such as human (**Supplementary Figure 6**, **7**, and **8**), mouse (**Figure 1**), and fruit fly (**Supplementary Figure 5**); 2) diverse subcellular compartments, including membrane-enclosed organelles (nucleus and mitochondrion) and open space (cell surface); 3) all commonly used proximity labeling tools, including peroxidases (APEX2 and HRP) and biotin ligases (BioID, TurboID, and miniTurbo); 4) different mass spectrometry quantification methods including label free (**Supplementary Figure 8**) and isobaric labeling (**Figure 1** and **Supplementary Figure 5**, **6**, and **7**). Moreover, PEELing features a plug-n’-play web service for a complete workflow (**Figure 1a**) including data visualization—all data panels shown here were directly from PEELing—as well as a Python package enabling advanced customization and integration with other bioinformatics tools. It thus provides a user-friendly yet versatile platform for exploring subcellular organization of the proteome.

## Acknowledgements

We thank members of the Scientific Computing Software team, the Open Science Software Initiative, and the Li laboratory at the Janelia Research Campus for support and advice on this project; L. Ding, M. Liu, L. Luo, C. McLaughlin, Y. Peng, M. Wang, Y. Wang, and Z. Zhang for feedback and comments on PEELing; and B. Smith and J. Lohmeyer for administrative assistance.

## Author Contributions

X.P., Data curation, Formal analysis, Software, Visualization, Methodology, Writing—review and editing; J.C., Data curation, Formal analysis, Software, Visualization, Methodology, Writing— review and editing; Z.J., Formal analysis, Writing—review and editing; S.H., Data curation, Methodology, Writing—review and editing, Resources, Funding acquisition; S.P., Software, Writing—review and editing, Supervision, Resources, Funding acquisition; J.L., Conceptualization, Data curation, Formal analysis, Visualization, Methodology, Writing— original draft, Supervision, Resources, Funding acquisition.

## Conflict of Interest

None declared.

## Funding

This work was supported by the Howard Hughes Medical Institute (to S.P. and J.L.), the National Key R&D Program of China (2023YFA1801300 and 2022YFA1304500 to S.H.), the Strategic Priority Research Program of the Chinese Academy of Sciences (XDB0570000 to S.H.), the National Natural Science Foundation of China (92368102 and 22377126 to S.H.), and the Shanghai Municipal Science and Technology Major Project (to S.H.).

## Data Availability

All datasets used for benchmarking are included as **Supplementary File 1–17**.

## Supplementary Materials

### Method Details

#### Data input

We note that PEELing does not handle raw mass spectral data or provide custom searches (e.g., labeling site identification or non-canonical translation analysis), which needs to be processed by peptide identification and quantification software such as MaxQuant (Cox and Mann, 2008), FragPipe (Kong *et al.*, 2017), Spectrum Mill (Broad Institute), or Proteome Discoverer (Thermo Fisher). The PEELing website and command line program accept tab-separated value (.tsv) files, which contain UniProt accession numbers in their first columns and enrichment indexes in the remaining columns (e.g., **Supplementary File 1–9**). PEELing uses UniProt accession numbers as protein identifiers and updates user input to the current UniProt version to help database searches. As benchmarked and discussed above, labelled-to-control ratio is the preferred enrichment index; however, users may choose other indexes according to their experimental design.

#### True positive and false positive references

We constructed true positive (TP) and false positive (FP) references using SwissProt-reviewed annotations. The PEELing web service automatically updates these references from the UniProt database (Consortium *et al.*, 2023), ensuring that they remain current and accurate.

Cell-surface TP reference (**Supplementary File 10**) is specified by the UniProt term: *((cc_scl_term:SL-0112) OR (cc_scl_term:SL-0243) OR (keyword:KW-0732) OR (cc_scl_term:SL-9906) OR (cc_scl_term:SL-9907)) AND (reviewed:true)*, which includes SwissProt-reviewed extracellular (SL-0112), secreted (SL-0243), signal peptide-containing (KW- 0732), type II transmembrane (SL-9906), or type III transmembrane (SL-9907) proteins. Cell-surface FP reference (**Supplementary File 12**) is specified by the UniProt term: *(((cc_scl_term:SL-0091) OR (cc_scl_term:SL-0173) OR (cc_scl_term:SL-0191)) AND (reviewed:true)) NOT (((cc_scl_term:SL-0112) OR (cc_scl_term:SL-0243) OR (keyword:KW- 0732) OR (cc_scl_term:SL-9906) OR (cc_scl_term:SL-9907)) AND (reviewed:true))*, including SwissProt-reviewed cytosolic (SL-0091), mitochondrial (SL-0173), and nuclear (SL-0191) proteins that do not express on the cell surface. Some cell-surface proteins, such as the Notch family proteins, are also localized in intracellular compartments and are not considered false positives, and are thus removed from the FP reference.

Nuclear TP reference (**Supplementary File 14**) is specified by the UniProt term:

*(cc_scl_term:SL-0191) AND (reviewed:true)*, which includes SwissProt-reviewed nuclear proteins (SL-0191). Nuclear FP reference (**Supplementary File 15**) is specified by the UniProt term: *(((cc_scl_term:SL-0091) OR (cc_scl_term:SL-0173) OR (cc_scl_term:SL-0112) OR (cc_scl_term:SL-0243) OR (keyword:KW-0732) OR (cc_scl_term:SL-9906) OR (cc_scl_term:SL- 9907)) AND (reviewed:true)) NOT ((cc_scl_term:SL-0191) AND (reviewed:true))*, including SwissProt-reviewed cytosolic (SL-0091), mitochondrial (SL-0173), and cell-surface (cell-surface TP term) proteins that do not express in the nucleus.

Mitochondrial TP reference (**Supplementary File 16**) is specified by the UniProt term: *(cc_scl_term:SL-0173) AND (reviewed:true)*, which includes SwissProt-reviewed mitochondrial proteins (SL-0173). Mitochondrial FP reference (**Supplementary File 17**) is specified by the UniProt term: *(((cc_scl_term:SL-0091) OR (cc_scl_term:SL-0191) OR (cc_scl_term:SL-0112) OR (cc_scl_term:SL-0243) OR (keyword:KW-0732) OR (cc_scl_term:SL-9906) OR (cc_scl_term:SL- 9907)) AND (reviewed:true)) NOT ((cc_scl_term:SL-0173) AND (reviewed:true))*, including SwissProt-reviewed cytosolic (SL-0091), nuclear (SL-0191), and cell-surface (cell-surface TP term) proteins that do not express in the mitochondrion.

For custom TP and FP references, PEELing requests two tab-separated value (.tsv) files from the user, each containing one column of UniProt accession numbers (e.g., **Supplementary File 10–17**). Despite the robustness of this algorithm and its tolerance to imperfect references (**Supplementary Figure 4**), constructing high-quality references is essential for proper data filtering and interpretation. The TP reference should contain proteins known to be in the designated cellular compartment while the FP reference should contain proteins that are, to one’s best knowledge, not in this compartment. Curated Swiss-Prot/UniProt and Gene Ontology Cellular Component (GOCC) databases, as well as relevant literature, provide resources for creating the TP and FP references. More TP and FP examples and their designing rules can be found in (Hung *et al.*, 2016) and (Cho *et al.*, 2020).

#### Reference-based data quality check and cutoff analysis

For each enrichment index (e.g., each labelled-to-control ratio), PEELing first ranks all proteins in descending order according to this index. For each protein on the ranked list, accumulated true positive count and false positive count above this ranking position are calculated to obtain true positive rate (TPR), false positive rate (FPR), and their difference (TPR–FPR) at this ranking position (visualized in **Figure 1d** and other plots of TPR, FPR, and TPR–FPR). A receiver operating characteristic (ROC) curve is produced accordingly (**Figure 1e** and other ROC curves).

For each enrichment index, the cutoff is set where TPR–FPR maximizes, representing the largest segregation of intended signal (TP) and unintended noise (FP). PEELing conducts cutoff analysis on all submitted enrichment indexes individually and, for the final proteome, retains only those proteins passing the cutoff of all or multiple indexes. Both the PEELing website and command line program offer the optional “Tolerance” setting, enabling users to control the stringency of the cutoff. By default, it is set to 0, meaning that a protein must pass the cutoff of all indexes to be included in the final proteome. If it is set to n, a protein can fail the cutoff in up to n indexes and still be included in the final proteome. Despite the flexibility, we recommend setting the tolerance value to a small number to better filter out contaminants.

#### Protein annotation

From the post-cutoff proteome, PEELing retrieves from UniProt and displays gene names, protein names, organisms, and protein lengths of the top 100 enriched proteins based on each index. For protein ontology, PEELing sends the post-cutoff proteome to the Panther server (Thomas *et al.*, 2022; Mi *et al.*, 2019) for over-representation analysis. The annotation datasets used include: GO slim cellular component, for protein localization; GO slim biological process, for protein function; and reactome pathway, for signaling pathway. Results are ranked in ascending order by false discovery rate (FDR). Top 10 terms are listed along with their FDRs.

## Supplementary Figures

**Supplementary Figure 1.**
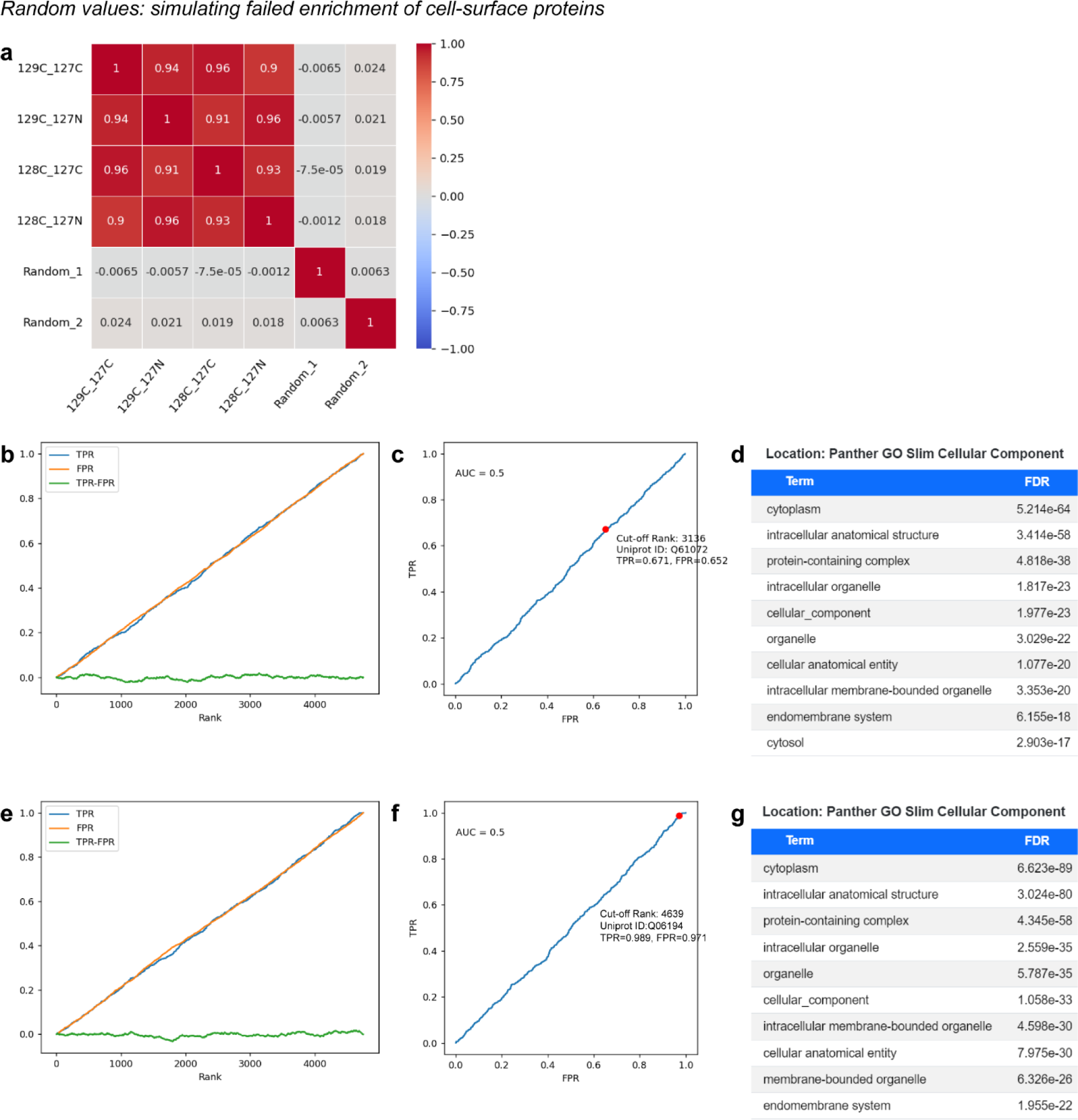
Testing PEELing with randomized data. To simulate failed enrichment, two sets of random values Random_1 and Random_2 were generated and added to the data of Purkinje cell cell-surface proteome (**Supplementary File 2**). (**a**) Correlation coefficients showing no correlation between the real and randomized data. (**b**,**e**) True positive rate (TPR, blue), false positive rate (FPR, orange), and their difference (TPR–FPR, green) plotted against Random_1 (**b**) and Random_2 (**e**) based ranking (x-axis). (**c**,**f**) Receiver operating characteristic (ROC) curves, based on Random_1 (**c**) and Random_2 (**f**), respectively. AUC, area under the curve. (**d**,**g**) Subcellular localization annotation did not enrich any cell surface related terms. **d**, Random_1; **g**, Random_2. FDR, false discovery rate.

**Supplementary Figure 2.**
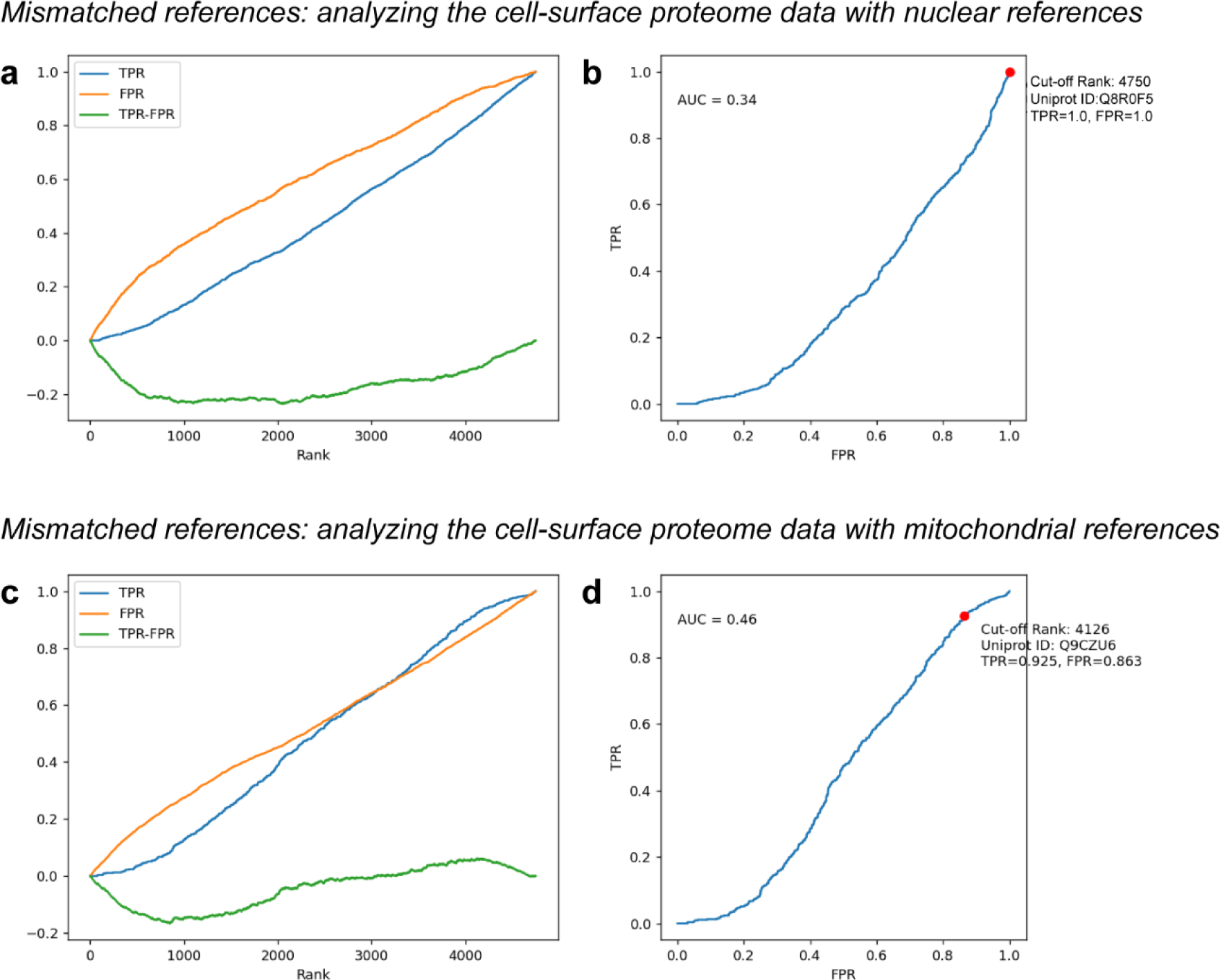
Testing PEELing with mismatched references. The cell-surface proteome of Purkinje cells (**Supplementary File 1**) was analyzed using mismatched references: nucleus (**a**,**b**; **Supplementary File 14**,**15**) and mitochondrion (**c**,**d**; **Supplementary File 16,17**). (**a**,**c**) True positive rate (TPR, blue), false positive rate (FPR, orange), and their difference (TPR–FPR, green) plotted against 129C:127N ratio-based ranking (x-axis). (**b**,**d**) Receiver operating characteristic (ROC) curves, based on ranking by 129C:127N. AUC, area under the curve.

**Supplementary Figure 3.**
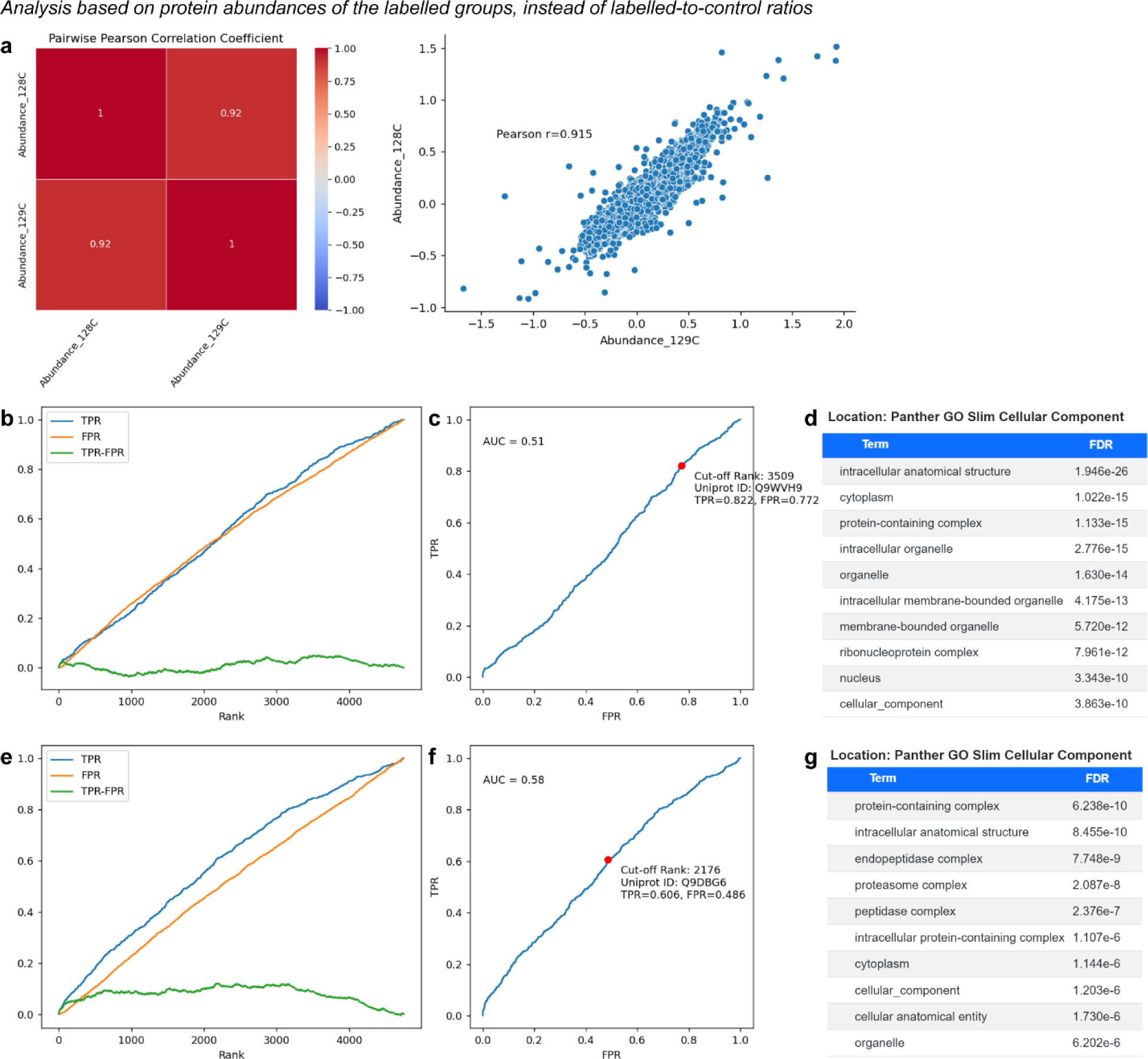
Abundance is not a reliable indicator of enrichment. Protein abundance data of cell-surface-labelled samples (TMT tags 128C and 129C) was obtained from the cell-surface proteome of Purkinje cells (Shuster *et al.*, 2022) (**Supplementary File 3**). (**a**) Correlation plots and coefficients. (**b**,**e**) True positive rate (TPR, blue), false positive rate (FPR, orange), and their difference (TPR–FPR, green) plotted against abundance-based ranking (x-axis). **b**, 128C; **e**, 129C. (**c**,**f**) Receiver operating characteristic (ROC) curves, based on ranking by abundance. **c**, 128C; **f**, 129C. AUC, area under the curve. (**d**,**g**) Subcellular localization of the abundance-ranked top 500 proteins. **d**, 128C; **g**, 129C. FDR, false discovery rate.

**Supplementary Figure 4.**
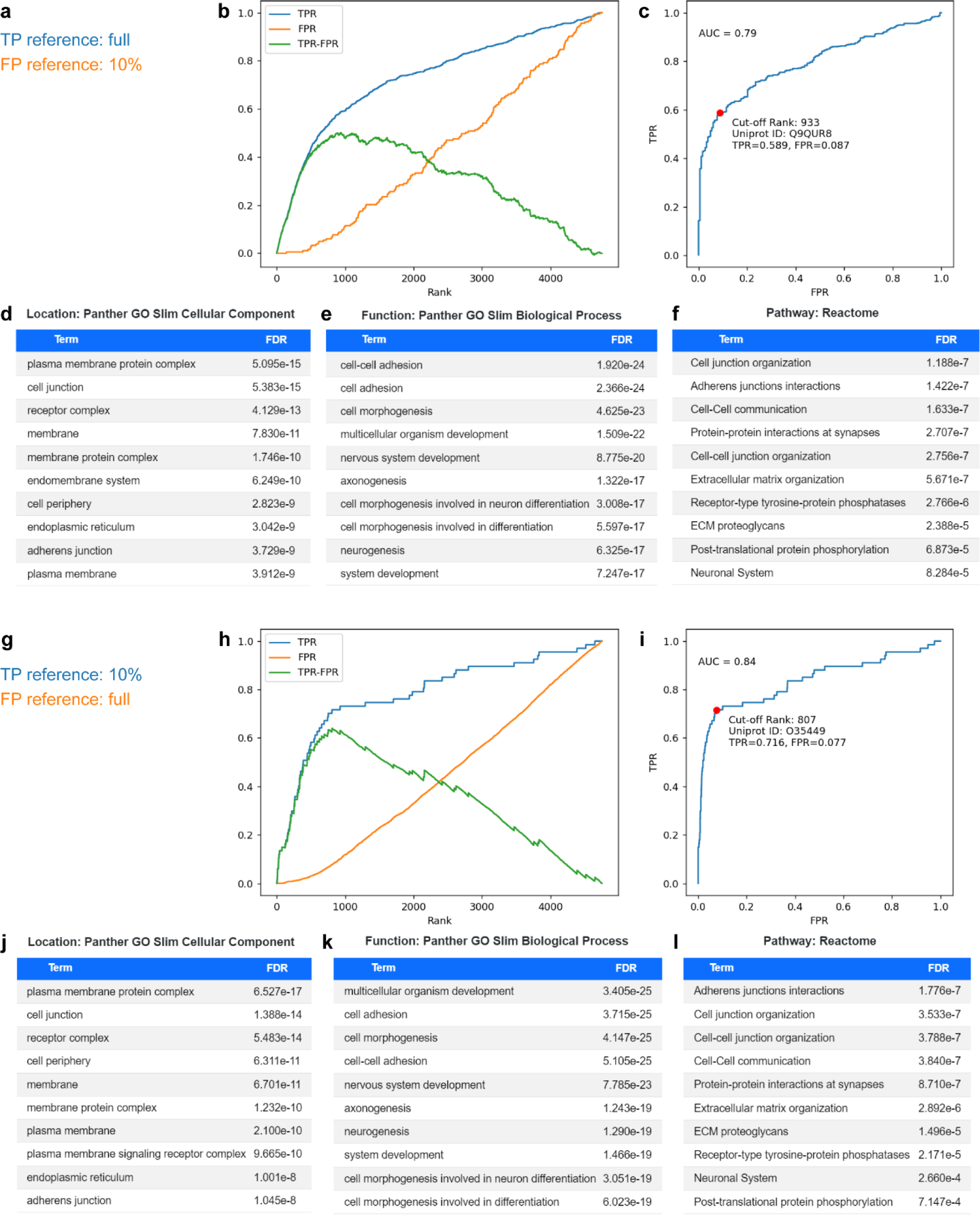
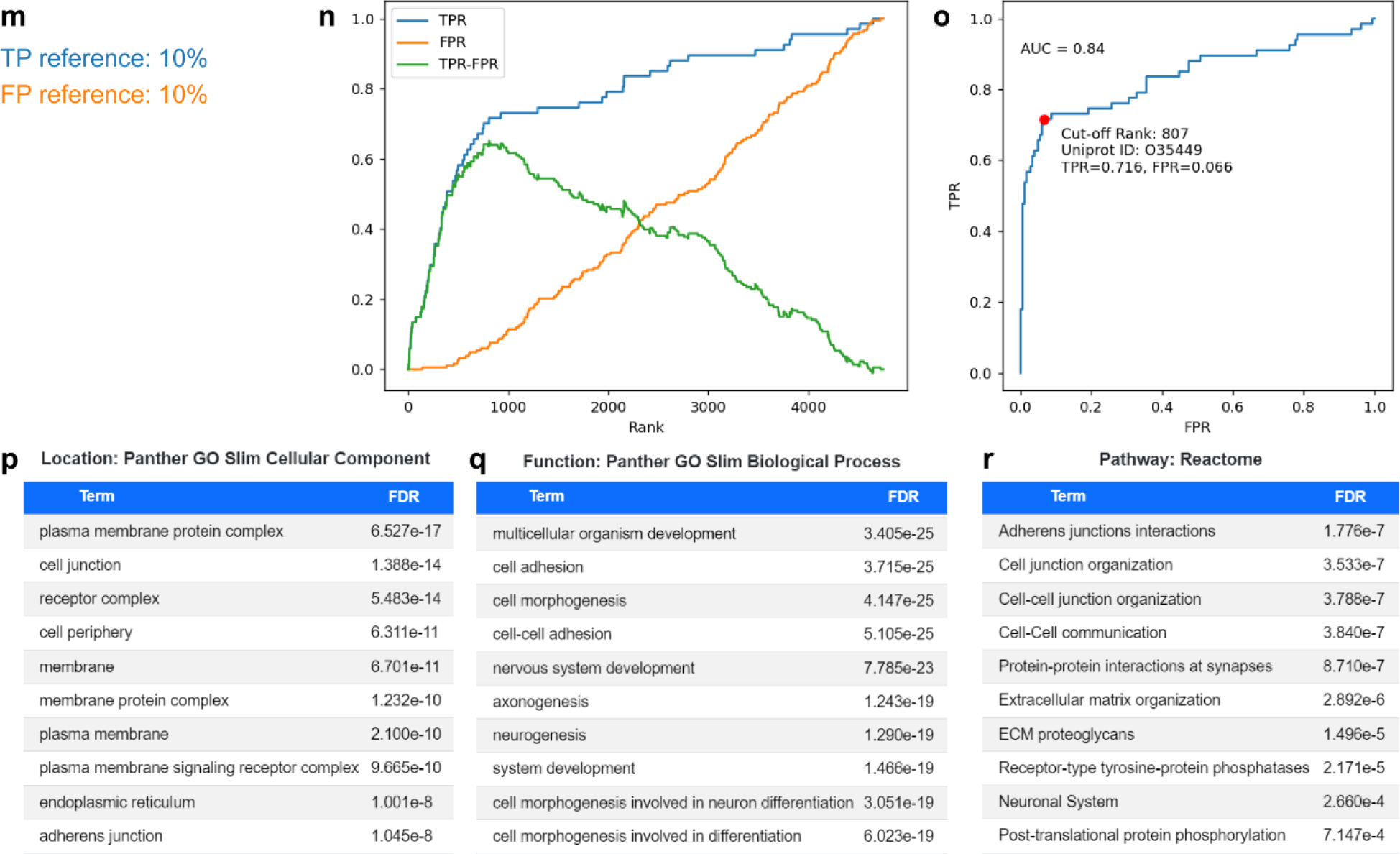
Testing PEELing with coverage-reduced references. We randomly removed 90% of genes from the cell-surface true-positive (TP) and false-positive (FP) references (**Supplementary File 10,12**) to produce truncated references with only 10% of the original coverage (**Supplementary File 11,13**) and then analyzed the cell-surface proteome of Purkinje cells (**Supplementary File 1**) using the truncated references (**m–r**) or pairing a full reference with a truncated one (**a–l**). (**b**,**h**,**n**) True positive rate (TPR, blue), false positive rate (FPR, orange), and their difference (TPR–FPR, green) plotted against 129C:127N ratio-based ranking (x-axis). (**c**,**i**,**o**) Receiver operating characteristic (ROC) curve, based on ranking by 129C:127N. Red dot, cutoff position. AUC, area under the curve. (**d–f**, **j–l**, **p–r**) Protein ontology analyses of post-cutoff proteomes for localization (**d**,**j**,**p**), function (**e**,**k**,**q**), and pathway (**f**,**l**,**r**). FDR, false discovery rate.

**Supplementary Figure 5.**
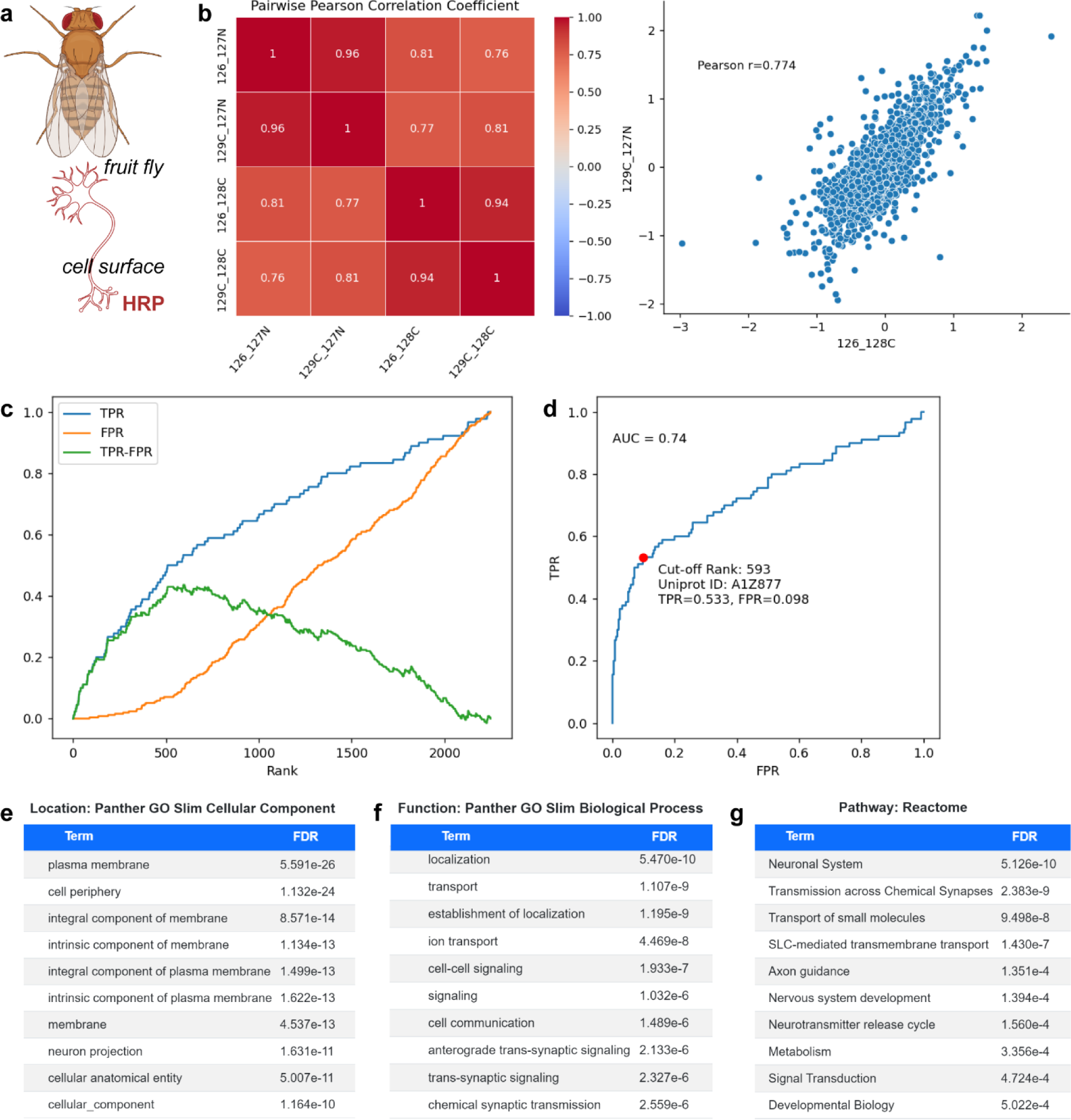
PEELing analysis of a *Drosophila* cell-surface proteome. (**a**) The cell-surface proteome of *Drosophila* mature olfactory projection neurons was profiled by horseradish peroxidase (HRP) based cell-surface proteomics (Li *et al.*, 2020) (**Supplementary File 4**). (**b**) Correlation plots and coefficients. (**c**) True positive rate (TPR, blue), false positive rate (FPR, orange), and their difference (TPR–FPR, green) plotted against 129C:127N ratio-based ranking (x-axis). (**d**) Receiver operating characteristic (ROC) curve, based on ranking by 129C:127N. Red dot, cutoff position. AUC, area under the curve. (**e–g**) Protein ontology analyses for localization (**e**), function (**f**), and pathway (**g**). FDR, false discovery rate.

**Supplementary Figure 6.**
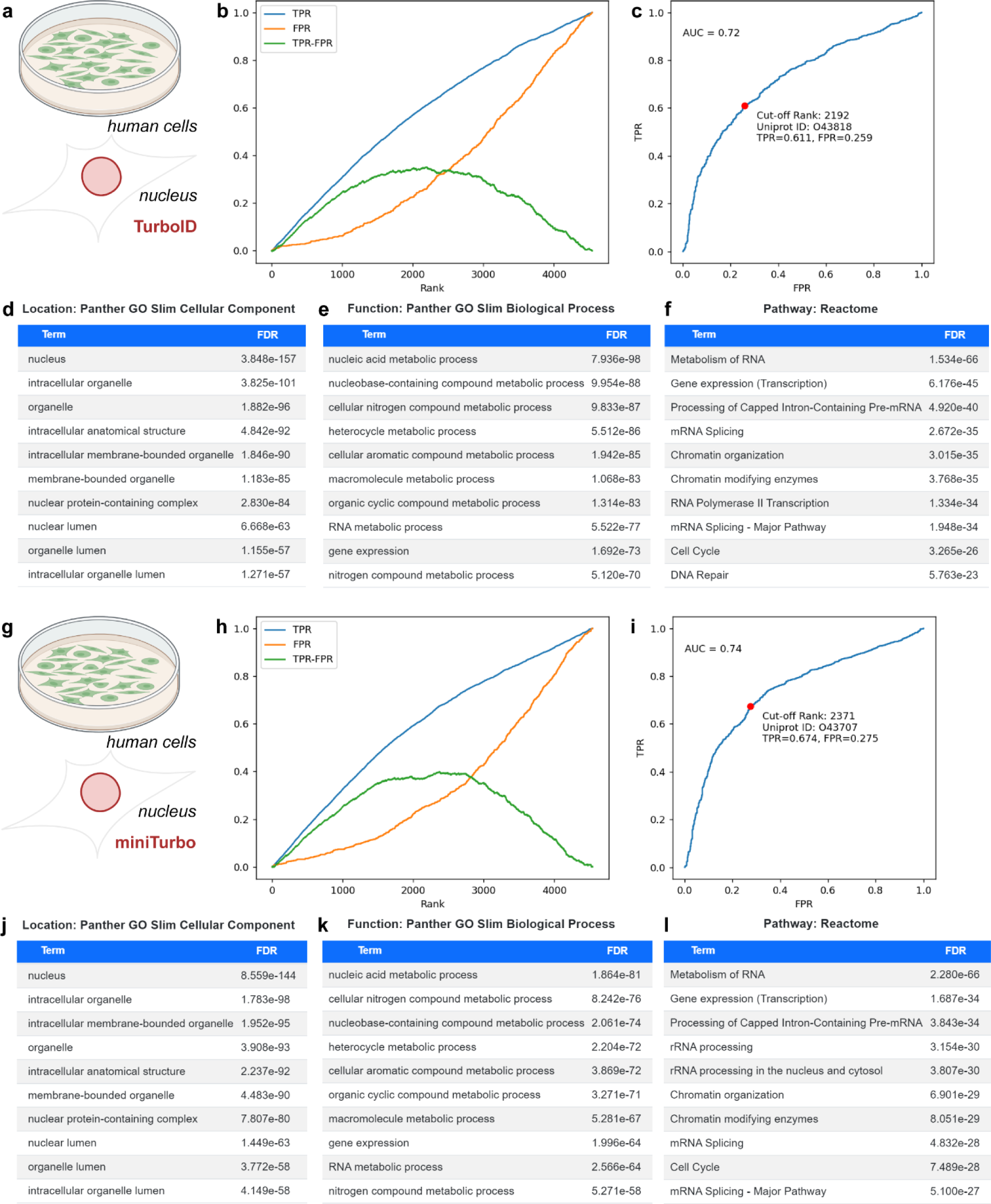
PEELing analysis of human nuclear proteomes. (**a–f**) Analysis of a nuclear proteome of human embryonic kidney 293 cells, profiled by (Branon *et al.*, 2018) using the biotin ligase TurboID (**Supplementary File 5**). (**g–l**) Analysis of a nuclear proteome of human embryonic kidney 293 cells, profiled by (Branon *et al.*, 2018) using the biotin ligase miniTurbo (**Supplementary File 6**). (**b**,**h**) True positive rate (TPR, blue), false positive rate (FPR, orange), and their difference (TPR–FPR, green) plotted against ratio-based ranking (x-axis). Ratio used: **b**, 129:126; **h**, 130:131. (**c**,**i**) Receiver operating characteristic (ROC) curve. Red dot, cutoff position. AUC, area under the curve. Ratio used: **c**, 129:126; **i**, 130:131. (**d–f**, **j–l**) Protein ontology analyses of post-cutoff proteomes for localization (**d**,**j**), function (**e**,**k**), and pathway (**f**,**l**). FDR, false discovery rate.

**Supplementary Figure 7.**
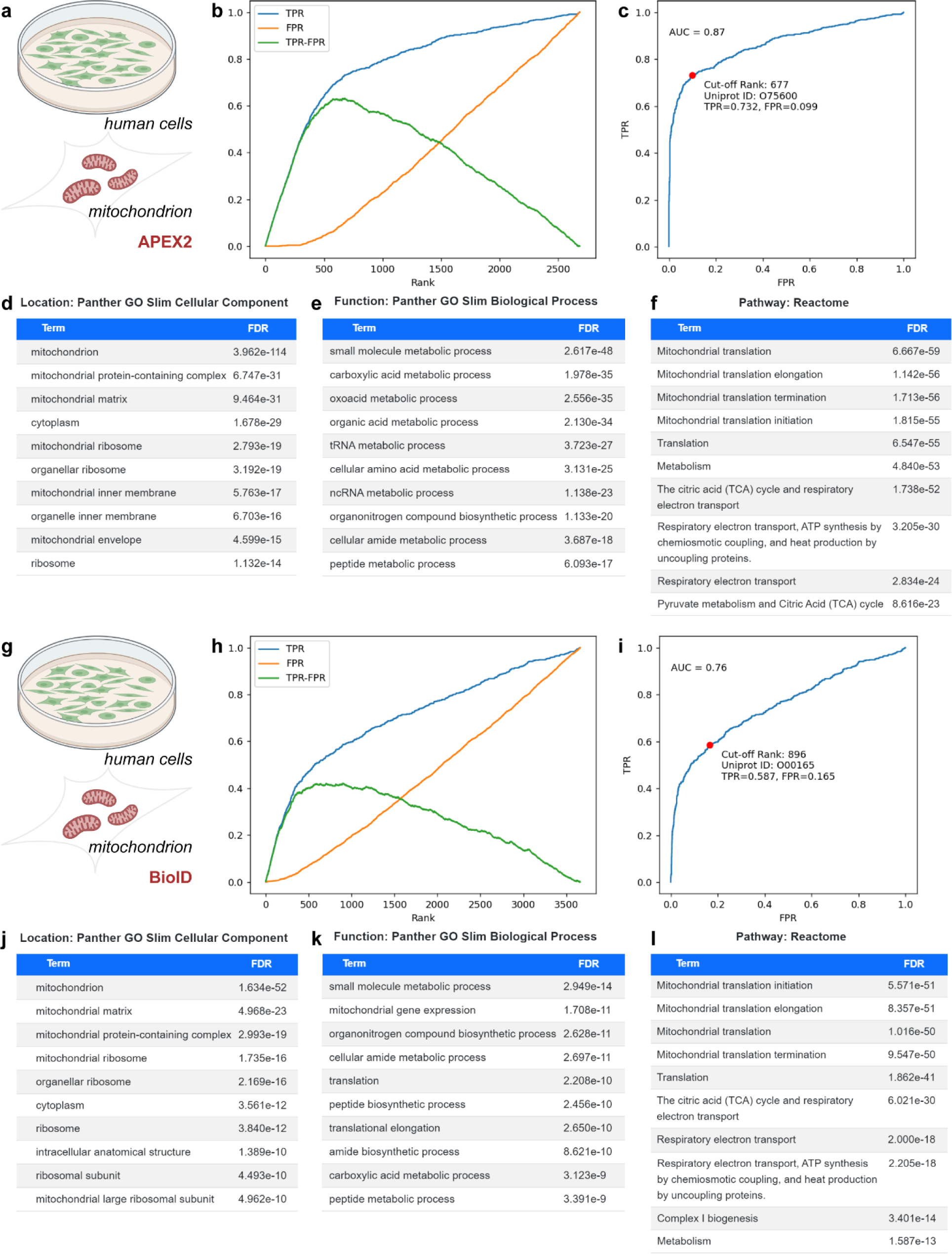
PEELing analysis of human mitochondrial proteomes. (**a–f**) Analysis of a mitochondrial nucleoid proteome of human embryonic kidney 293 cells, profiled by (Han *et al.*, 2017) using the peroxidase APEX2 (**Supplementary File 7**). (**g–l**) Analysis of a mitochondrial matrix proteome of human embryonic kidney 293 cells, profiled by (Branon *et al.*, 2018) using the biotin ligase BioID (**Supplementary File 8**). (**b**,**h**) True positive rate (TPR, blue), false positive rate (FPR, orange), and their difference (TPR–FPR, green) plotted against ratio-based ranking (x-axis). Ratio used: **b**, 126:128; **h**, 126:127. (**c**,**i**) Receiver operating characteristic (ROC) curve. Red dot, cutoff position. AUC, area under the curve. Ratio used: **c**, 126:128; **i**, 126:127. (**d–f**, **j–l**) Protein ontology analyses of post-cutoff proteomes for localization (**d**,**j**), function (**e**,**k**), and pathway (**f**,**l**). FDR, false discovery rate.

**Supplementary Figure 8.**
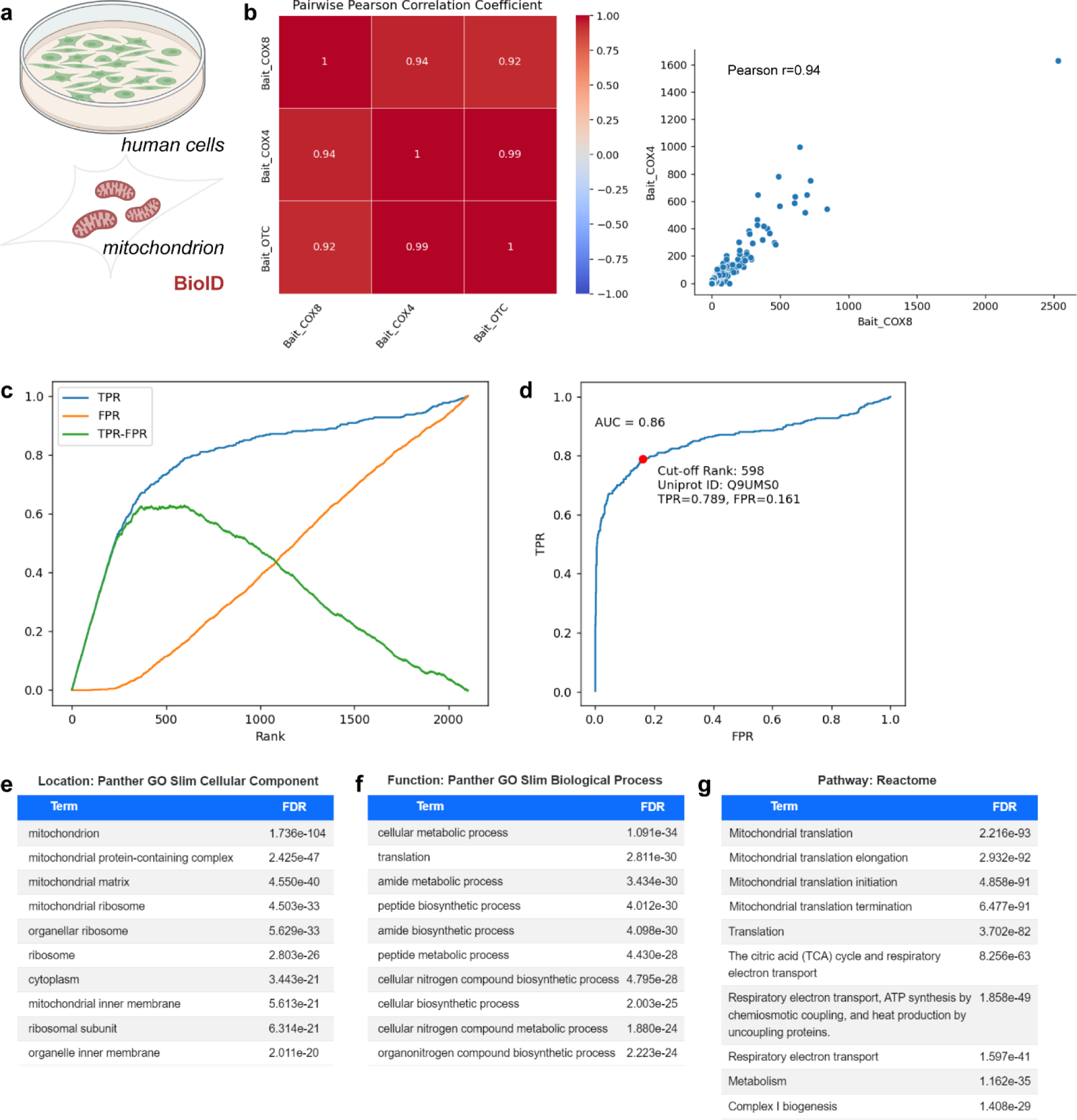
PEELing analysis of a label-free quantification mass spectrometry dataset. (a) Analysis of a mitochondrial matrix proteome of human embryonic kidney 293 cells, profiled by (Antonicka *et al.*, 2020) using the biotin ligase BioID fused to three different bait proteins COX4, COX8, and OTC and label-free quantification mass spectrometry (**Supplementary File 9**). (b) Correlation plots and coefficients. (**c**) True positive rate (TPR, blue), false positive rate (FPR, orange), and their difference (TPR–FPR, green) plotted against the labelled-to-control ratio-based ranking of the COX4 bait group (x-axis). (**d**) Receiver operating characteristic (ROC) curve, based on the labelled-to-control ratio-based ranking of the COX4 bait group. Red dot, cutoff position. AUC, area under the curve. (**e–g**) Protein ontology analyses for localization (**e**), function (**f**), and pathway (**g**). FDR, false discovery rate.

## Supplementary Files

**Supplementary File 1.** A cell-surface proteome of mouse developing Purkinje cells (Shuster *et al.*, 2022). The authors used horseradish peroxidase (HRP) based cell-surface biotinylation and tandem mass tag (TMT) (Thompson *et al.*, 2003) based quantitative mass spectrometry. TMT tags 129C and 128C were used for labelled samples while 127C and 127N were used for non-labelled controls, producing four labelled-to-control ratios (129C:127C, 129C:127N, 128C:127C, and 128C:127N). Ratios are normalized and log2-transformed.

**Supplementary File 2.** Two sets of random values, Random_1 (6^th^ column) and Random_2 (7^th^ column), were generated and added to the data of mouse Purkinje cell cell-surface proteome (Shuster *et al.*, 2022) (1^st^ to 5^th^ columns; from **Supplementary File 1**).

**Supplementary File 3.** Protein abundance data of the cell-surface proteome of mouse developing Purkinje cells (Shuster *et al.*, 2022), derived from the same mass spectrometry data as **Supplementary File 1**. The authors used horseradish peroxidase (HRP) based cell-surface biotinylation and tandem mass tag (TMT) (Thompson *et al.*, 2003) based quantitative mass spectrometry. TMT tags 129C and 128C were used for labelled samples. Abundance values are normalized and log2-transformed.

**Supplementary File 4.** A cell-surface proteome of *Drosophila* mature olfactory projection neurons (Li *et al.*, 2020). The authors used horseradish peroxidase (HRP) based cell-surface biotinylation and tandem mass tag (TMT) (Thompson *et al.*, 2003) based quantitative mass spectrometry. TMT tags 126 and 129C were used for labelled samples while 127N and 128C were used for non-labelled controls, producing four labelled-to-control ratios (126:127N, 129C:127N, 126:128C, and 129C:128C). Ratios are normalized and log2-transformed.

**Supplementary File 5.** A nuclear proteome of human embryonic kidney 293 cells (Branon *et al.*, 2018). The authors used the biotin ligase TurboID for spatially-resolved biotinylation in the nucleus and tandem mass tag (TMT) (Thompson *et al.*, 2003) for quantitative mass spectrometry. Two labelled-to-control ratios were provided: TMT tag 128:129 from Replicate 1 and 129:126 from Replicate 2. Ratios are normalized and log2-transformed.

**Supplementary File 6.** A nuclear proteome of human embryonic kidney 293 cells (Branon *et al.*, 2018). The authors used the biotin ligase miniTurbo for spatially-resolved biotinylation in the nucleus and tandem mass tag (TMT) (Thompson *et al.*, 2003) for quantitative mass spectrometry. Two labelled-to-control ratios were provided: TMT tag 130:131 from Replicate 1 and 127:126 from Replicate 2. Ratios are normalized and log2-transformed.

**Supplementary File 7.** A mitochondrial nucleoid proteome of human embryonic kidney 293 cells (Han *et al.*, 2017). The authors used the peroxidase APEX2 for spatially-resolved biotinylation in the mitochondrion and tandem mass tag (TMT) (Thompson *et al.*, 2003) for quantitative mass spectrometry. Two labelled-to-control ratios were provided: TMT tag 126:128 from Replicate 1 and 129:131 from Replicate 2. Ratios are normalized and log2-transformed.

**Supplementary File 8.** A mitochondrial matrix proteome of human embryonic kidney 293 cells (Branon *et al.*, 2018). The authors used the biotin ligase BioID for spatially-resolved biotinylation in the mitochondrion and tandem mass tag (TMT) (Thompson *et al.*, 2003) for quantitative mass spectrometry. Two labelled-to-control ratios were provided: TMT tag 126:127 from Replicate 1 and 130:131 from Replicate 2. Ratios are normalized and log2-transformed.

**Supplementary File 9.** A mitochondrial matrix proteome of human embryonic kidney 293 cells (Antonicka *et al.*, 2020). The authors used the biotin ligase BioID fused to three different bait proteins COX4, COX8, and OTC for spatially-resolved biotinylation in the mitochondrion and label-free quantification mass spectrometry. One labelled-to-control ratio was provided for each bait.

**Supplementary File 10.** True positive (TP) reference for the cell surface, as of July 2, 2023.

**Supplementary File 11.** 10%-coverage TP reference for the cell surface, produced by randomly removing 90% of genes from

**Supplementary File 12.** False positive (FP) reference for the cell surface, as of July 2, 2023.

**Supplementary File 13.** 10%-coverage FP reference for the cell surface, produced by randomly removing 90% of genes from

**Supplementary File 14.** True positive (TP) reference for the nucleus, as of July 2, 2023.

**Supplementary File 15.** False positive (FP) reference for the nucleus, as of July 2, 2023.

**Supplementary File 16.** True positive (TP) reference for the mitochondrion, as of July 2, 2023.

**Supplementary File 17.** False positive (FP) reference for the mitochondrion, as of July 2, 2023.

## Notes

https://peeling.janelia.org/

https://github.com/JaneliaSciComp/PEELing

### Competing Interest Statement

The authors have declared no competing interest.

### Summary of Updates

We expanded the scope and utility of PEELing and benchmarked it with more diverse datasets.

